# How plants minimize somatic evolution

**DOI:** 10.1101/2021.02.13.431063

**Authors:** Máté Kiss, Gergely J. Szöllősi, Imre Derényi

## Abstract

A remarkable property of plants is their ability to accumulate mutations at a very slow pace despite their potentially long lifespans, during which they continually form buds, each with the potential to become a new branch. Because replication errors in cell division represent an unavoidable source of mutations, minimizing mutation accumulation requires the minimization of cell divisions. Here we show that there exists a well defined theoretical minimum for the branching cost, defined as the number of cell divisions necessary for the creation of each branch. Most importantly, we also show that this theoretical minimum can be closely approached by a simple pattern of cell divisions in the meristematic tissue of apical buds during the generation of novel buds. Both the optimal pattern of cell divisions and the associated branching cost are consistent with recent experimental data, suggesting that plant evolution has led to the discovery of this mechanism.

## INTRODUCTION

Trees can live for centuries or even millennia [1–3], reaching tens of (sometimes more than a hundred) meters in height [4] and hundreds of cubic meters in volume [5]. Their growth requires mechanisms with which they can protect their genomes from excessive mutation accumulation both to avoid tissue deterioration and to maintain the genetic fidelity of their gametes. Such mechanisms must involve the minimization of the number of cell divisions leading from the zygote to the gametes, because replication errors (in the range of 10^−9^ single nucleotide mutations (SNMs) per base pair (bp) per cell division) are an inevitable source of mutations.

Multicellular animals usually possess a germline, which is segregated early on during development. It is unclear if plants also have a segregated germline [6], although true germlines are possibly present in tissues of certain species [7], evidence suggests the existence of a so-called functional germline in most plants, which is responsible for the production of both somatic and reproductive cells [6]. As an example, mutations in cells taken from branches of an oak can also be found in the acorns grown from the same branches [8].

Irrespective of having a true or functional germline, plants face the problem that during their growth axillary buds (green shapes in Fig. 1, harboring undifferentiated meristematic tissues [9]) at the leaf axils are continually created, each having the potential to turn into an apical (terminal) bud (yellow shapes in Fig. 1) and initiate a new branch. Bud creation, thus, leads to cell lineage bifurcation, and necessitates cell divisions.

**FIG. 1.**
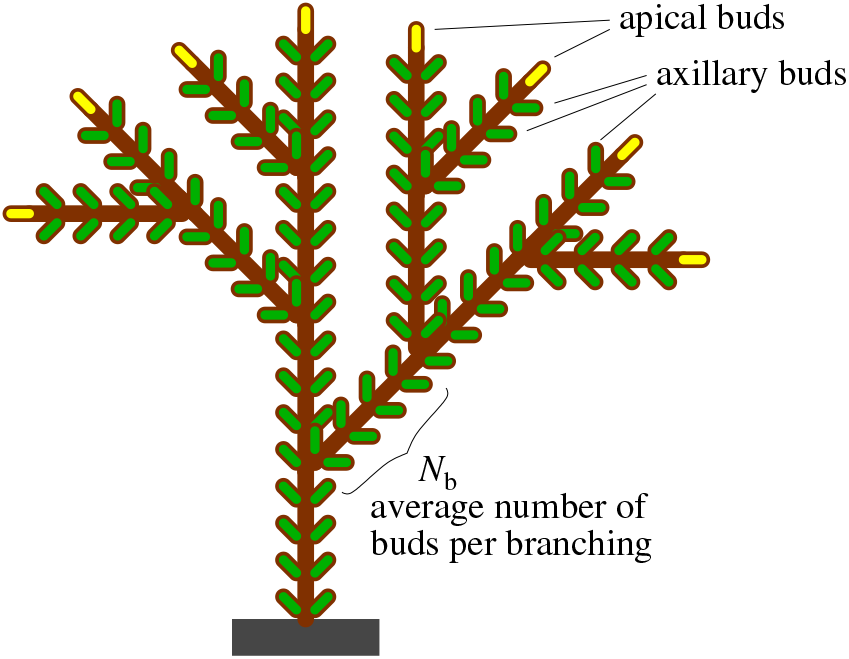
A schematic illustration of a plant with multiple branches. Apical (terminal) and axillary buds are represented by yellow and green shapes, respectively. The average number of axillary buds per branching is denoted by *N*_b_. The branching order of this plant is 3, corresponding to 2^3^ = 8 terminal branches and apical buds.

Schmid-Siegert et al. [10] have recently found a mere 17 single nucleotide variations (SNVs) between leaves from two distant branches of a 234-year-old oak. Even after correcting for missed variations, they estimate the total number of SNVs to be no more than 47. This corresponds to the accumulation of no more than 24 SNMs along either branch from the moment of their separation when the tree was young. Using a conservative estimate of 0.3 × 10^−9^ 1/bp for the mutation rate and a size of 0.8 × 10^9^ bp for the oak genome, the number of SNMs per cell division is about 0.24, indicating that the number of cell divisions leading from the moment of branch separation to the tip of either branch is no more than 24*/*0.24 = 100. This is a remarkably low number, considering the fact that of the order of 1000 axillary meristems must have been created along these 20-meter-long branches.

Put differently, an average branching order (number of branching events from the trunk to the tips) of about 20 (being responsible for the generation of a million terminal branches), corresponds to no more than *c* ≈ 100*/*20 = 5 cell divisions per branching (referred to as *branching cost* from now on), even though there are as many as *N*_b_ ≈ 1000*/*20 = 50 axillary buds generated per branching. The branching cost of *c* ≈ 5 is also in line with the finding [11] that the number of cell divisions between the stem cells of the parent meristem and an axillary meristem in tomato is *c ≈* 9, and in *Arabidopsis* it is *c ≈* 7.

This remarkably low branching cost (despite the potentially large number of axillary buds generated per branching) requires explanation. The questions arise what the theoretical minimum value of the branching cost (*c*^min^) is, and how it can be approached by a cell divisional mechanism in the meristems.

## RESULTS

Both the apical and the axillary buds (yellow and green shapes in Fig. 1, respectively) contain a small tissue, called the meristem, the central zone of which consists of stem cells, surrounded by several layers of increasingly differentiated cells [6]. The axillary buds have the potential to turn into apical buds, thereby initiating the growth of a new branch. However, only a fraction of 1*/N*_b_ of them start new branches during the lifetime of a plant, where *N*_b_ denotes the average number of axillary buds per branching (Fig. 1).

Following the fates of the stem cells between two branching points, the number of stem cell divisions can be determined. Assuming the simplest (designated as “linear”) strategy for meristem creation (left panel of Fig. 2) the stem cells of the apical meristem (yellow line) divides into two whenever a new meristem is created, and half of the daughter stem cells (green lines) go into the new meristem. After the creation of *N*_b_ axillary meristems both the apical and the axillary meristems have gone through *N*_b_ cell divisions, resulting in *c* = *N*_b_ for the branching cost.

**FIG. 2.**
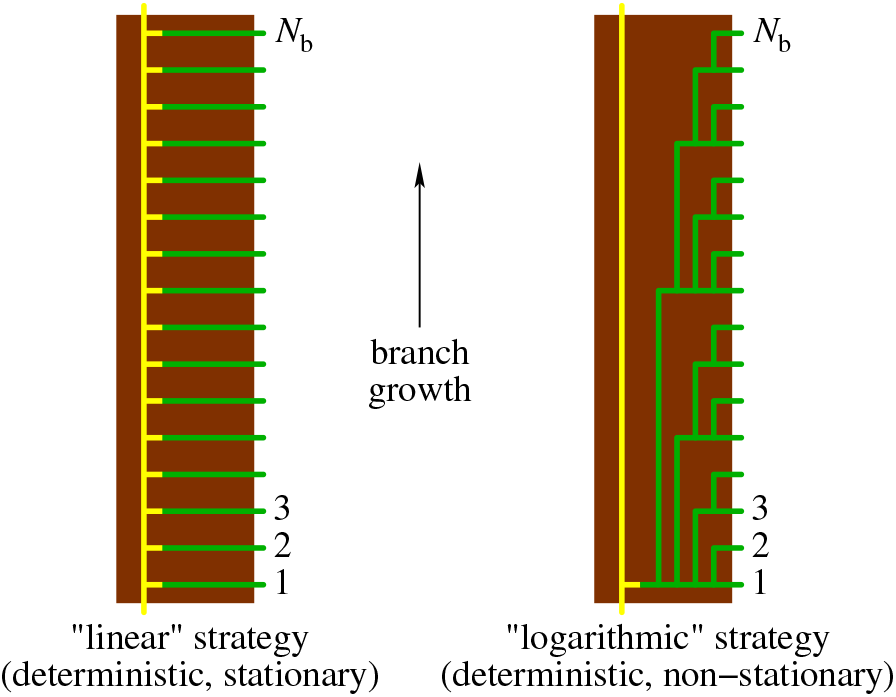
Stem cell fates in two strategies for meristem creation in a branch segment between two branching points. Left panel: “linear” strategy, in which the apical stem cells (yellow line) divide (green lines) whenever an axillary meristem is created. Right panel: “logarithmic” strategy, in which the apical stem cells (yellow line) divide only once, and the stem cells of the axillary meristems are generated by several rounds of cell divisions along a perfect binary tree (green lines).

The high branching cost of this strategy comes from the large number of divisions of the stem cells of the apical meristem. This high demand on the apical meristem can be minimized (right panel of Fig. 2) if its stem cells divide only once, and half of the daughter cells continue their way without further cell divisions (yellow line), while the other half go through an additional log_2_(*N*_b_) rounds of cell divisions to generate all the stem cells of the *N*_b_ axillary meristems along a perfect binary tree (green lines). Note that the slightly (only logarithmically) elevated number of cell divisions by the axillary meristematic stem cells is a small price to be paid for having the ability to generate a large number of meristems, out of which only a small fraction will initiate a new branch. Because on average only one of the *N*_b_ axillary buds turns into a branch, the average branching cost of this elaborate (designated as “logarithmic”) strategy is the arithmetical mean of the number of stem cell divisions along both the apical line (1) and one of the axillary lines (1 + log_2_(*N*_b_):

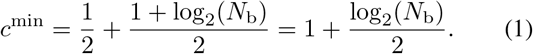

This is the theoretical minimum value of the average branching cost, as any cell divisions delegated from the axillary lines to the apical line will decrease the number of divisions of the axillary stem cells less than it will increase the number of divisions of the apical cells.

Using the estimation of *N*_b_ *≈* 50 for oak, the minimum branching cost is *c*^min^ *≈* 3.8, which is not much smaller than the estimated value of *c ≈* 5. This suggests oaks (and probably most plants) have evolved a mechanism with which the theoretical minimum branching cost can be approached.

The difficulty with the implementation of the “logarithmic” strategy is that it is both deterministic and non-stationary. Its deterministic feature means that the divisions of the stem cells follow a preset order, which requires precise bookkeeping, and does not tolerate errors. After running out of the axillary stem cells or initiating a new branch, the exact same sequence of events is repeated. The non-stationary nature of the strategy means that the generation of each axillary meristem within a sequence is different.

Our goal is to find a biologically feasible mechanism that is *stochastic* and *stationary*, such that each meristem is generated in the same fashion, which requires no bookkeeping (and no memory). To this end we introduce a general framework (Fig. 3), in which the central zone of a meristem consists of *n* cells (stem cells) that are arranged into a series of *n* levels (enumerated from 0 to *n* − 1) according to their distance from the center. In each time step (measured in units of meristem creation) one of the cells of an apical meristem at level *i* is selected for division with probability *p_i_* from a normalized probability distribution (i.e., 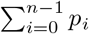), after which all the cells at levels higher than *i* are pushed upward by one level (i.e., *j → j* + 1 if *j > i*), and one of the daughter cells is placed into level *i* + 1, while the other one is retained at level *i*. An axillary meristem (the green one) is created by separating the cell that has been pushed to the extra level *n* of the apical meristem, making it the founder cell of the new meristem, and letting it go through a number of cell divisions
to build up the new tissue. The optimal method to do this is dividing according to a perfect binary tree, which provides a uniform divisional cost of

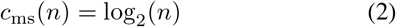

to each cell of the new meristem. The motivation behind this framework is that hierarchically organized tissues can efficiently reduce somatic evolution [12].

**FIG. 3.**
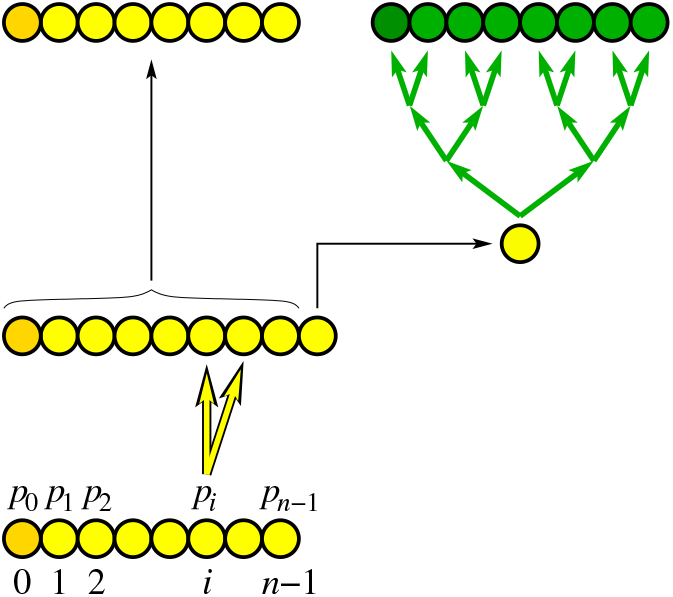
General framework for the mechanism of the generation of an axillary meristem. The central zone of meristems consists of *n* cells, organized into *n* levels. A new axillary meristem (green one) is created from an apical meristem (yellow one) by a series of cell divisions as depicted and described in detail in the main text. The cells at level 0 are marked by a darker shade.

After an axillary meristem turns into an apical one (at time set to *t* = 0) the average number of divisions *D_i_*(*t*) of the cell at level *i* starts to increase (from *D_i_*(0) = 0) due to cell divisions taking place in the meristem. At each timestep *D_i_*(*t*) is updated by the following equation:

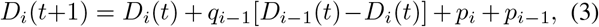

where the second term describes that cell from level *i* − 1 are pushed upward by probability

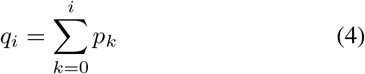

(with *q*_−1_ = 0), and the last two terms describe that the divisional number increases by one if a cell division takes place either at level *i* or *i* − 1 (with probabilities *p_i_* or *p_i_*_−1_, respectively). This equation can be used even for the cell at level *n*, which is the founder cell of the new meristem.

The average divisional cost per branching (*c*) can be calculated in the following way. By selecting a random path from the root of the tree to one of its terminal branches, at each branching point we have 50% chance to remain on the original branch and 50% chance to switch to the newly created branch. This means that for a random path switching to a new meristem occurs after 2*N*_b_ axillary buds on average. Therefore, the average divisional cost of the founder cell of the new branch where switching occurs can be calculated as

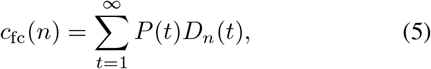

where

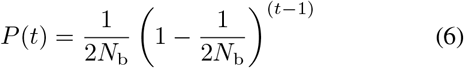

is the geometric distribution with success probability 1*/*(2*N*_b_).

Thus, the average divisional cost (number of cell divisions) from one switching to the next one is the sum of *c*_fc_(*n*) for generating the founder cell of the new meristem and *c*_ms_(*n*) for building up the new meristem. As this cost incurs after the generation of an average of 2*N*_b_ axillary buds, but between to branching points there are an average of only *N*_b_ axillary buds, the average branching cost is given by

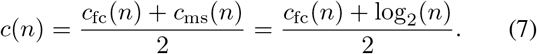

For any given *N*_b_ the independent parameters of this mechanism are the number of levels (*n*) and the divisional probabilities (*p_i_*) of the levels. What we are interested in is the optimal values of these parameters that minimize the average branching cost. Because Eq. (3) cannot be solved analytically, we have performed numerical optimization (for more details see the Methods section). To explore the effect of the size of the meristem, first we fixed the number of levels. The optimal probability distribution of cell divisions is demonstrated in Fig. 4 for *N*_b_ = 20 and at four different values of the number of levels (cells) of the meristem.

**FIG. 4.**
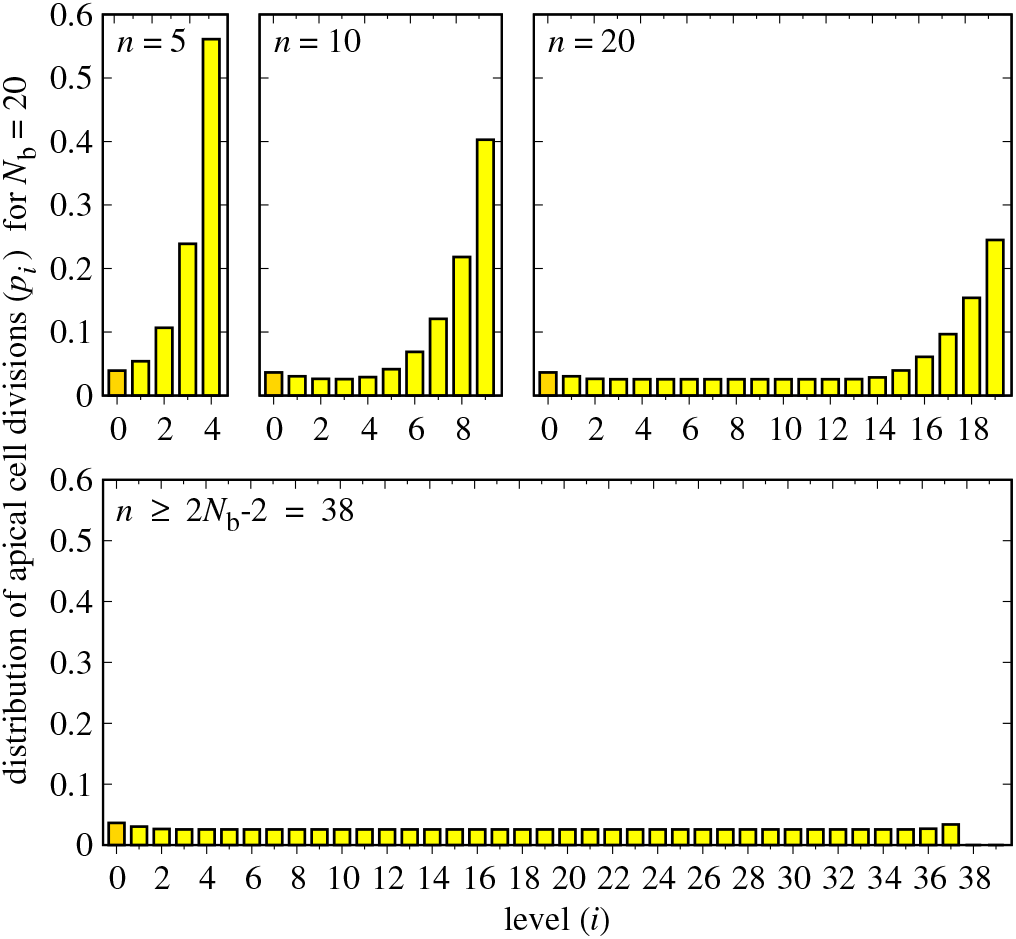
The optimal probability distribution of apical cell divisions for *N*_b_ = 20. In the four subfigures the number of levels of the meristem are set to *n* = 5, 10, 20, 40.

The general features of the optimal probability distribution are the following. For small values of *n* the divisional probability progressively increases towards higher levels. For increasing *n* the probabilities roll out like a carpet: they converge to constant (and almost uniform) values, except for the last few probabilities, which carry the excess probability. The “rolling out” stops at *n* = 2*N*_b_ *—* 2, and for larger values of *n* the first 2*N*_b_ − 2 probabilities remain unchanged and non-zero, while all the rest become zero (i.e., cells only travel through these levels, and never divide). These non-zero probabilities have almost a uniform value of 1*/*(2*N*_b_ − 1), except near the two ends, where they are slightly larger. Note that even with a uniform distribution of 1*/*(2*N*_b_ − 2), the average branching cost is almost the same.

For *N*_b_ = 20 the average branching cost *c*^opt^(*n*) for the optimal probability distribution is plotted in Fig. 5 as function of the number of levels (*n*) of the meristem. What is remarkable is that for a wide range of the number of levels the average branching cost is very close to the theoretical minimum (with a deviation of smaller than one), which proves that a stochastic and stationary mechanism of meristem generation is sufficient to bring down the branching cost near to the absolute theoretical minimum.

**FIG. 5.**
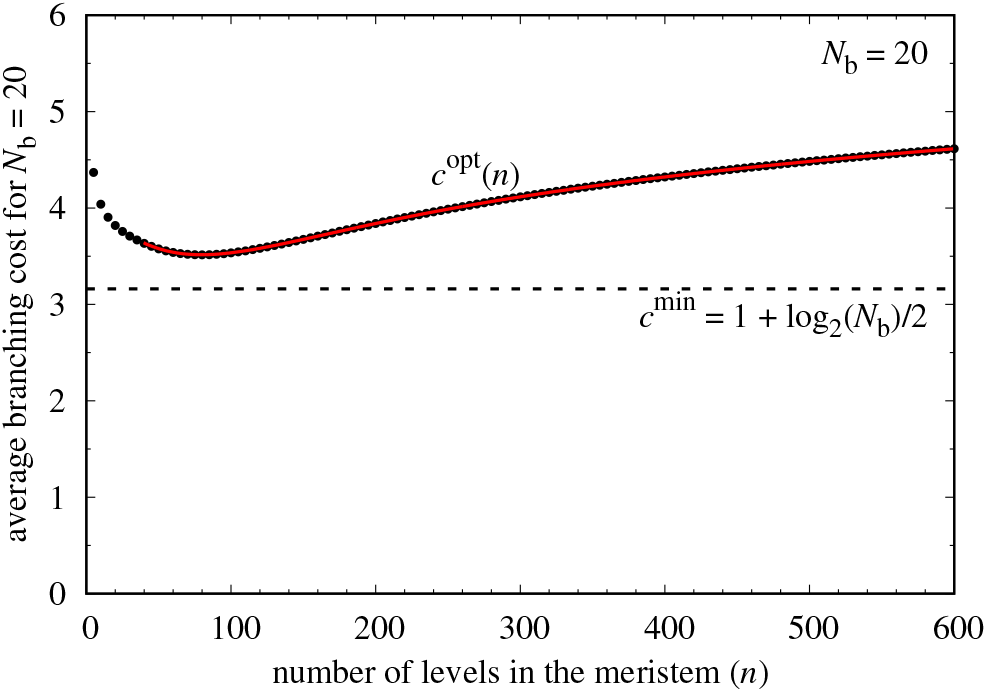
The branching cost as a function of *n* for *N*_b_ = 20. Dots are the numerically optimized values for fixed values of *n*, the red line is the approximative formula given by Eq. (10), and the dashed line is the theoretical minimum.

Due to the property of the optimal distribution that the probabilities are exactly zero for all the levels above the first 2*N*_b_ *−* 2 ones, the following relationship can be written up for *n ≥* 2*N*_b_ *−* 2 and *m ≥* 2*N*_b_ *−* 2:

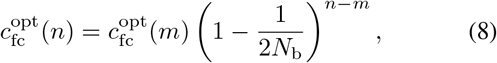

Interestingly, we find that 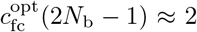 irrespective of the value of *N*_b_. Therefore, for *n ≥* 2*N*_b_ *−* 2

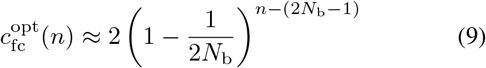

and

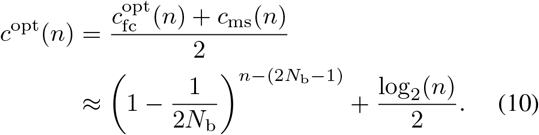

This approximation is also plotted in Fig. 5 with a red line, showing a very good agreement with the numerically obtained data.

The *c*^opt^(*n*) function has a minimum at around *n*^opt^ ≈ 1.02 × 4*N*_b_ ≈ 4*N*_b_. This minimal value, simply denoted by *c*^opt^, is plotted in Fig. 6. Plugging *n*^opt^ ≈ 4*N*_b_ into Eq. (10) results in

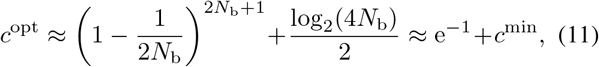

which, as can be seen in Fig. 6, is also in very good agreement with the numerical results. Thus, for the optimal probability distribution of the apical cell divisions, and for the optimal meristem size, the average branching cost is larger than the theoretical minimum by only about e^−1^ *≈* 0.368.

**FIG. 6.**
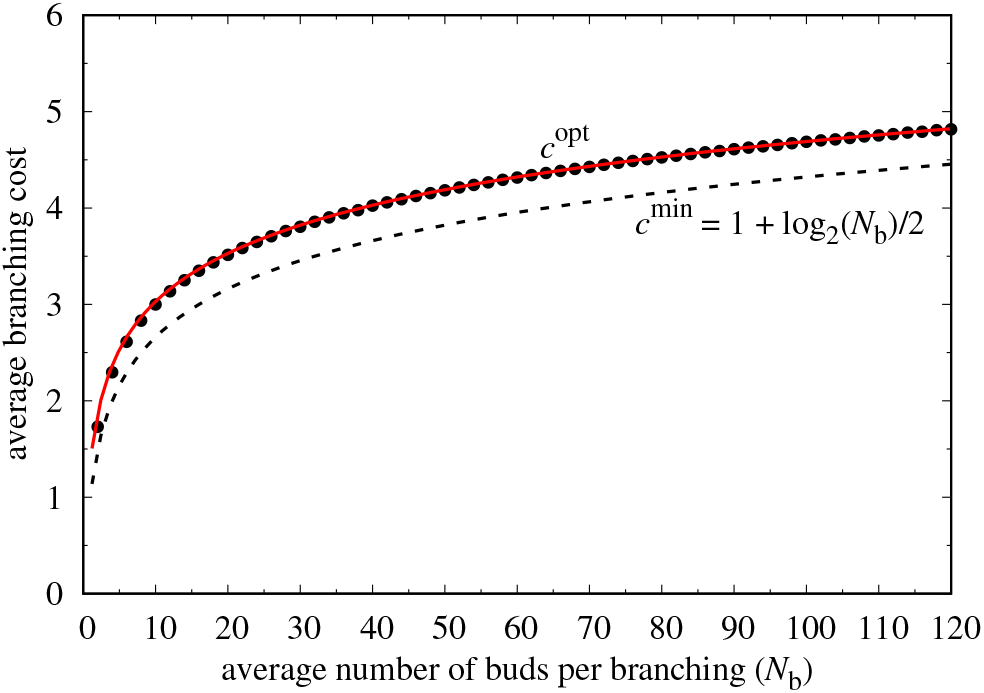
The branching costs as a function of *N*_b_. Dots are the numerically optimized values, the red line is the approximative formula given by Eq. (11), and the dashed line is the theoretical minimum.

## DISCUSSION

Our results demonstrate that by employing the mechanism described above, it is possible to achieve, in a stochastic manner, a number of cell divisions per branching close to the theoretical minimum. Due to the flat shape of the branching cost *c*^opt^(*n*) as a function of the number of levels, *n*, almost equally low branching costs can be achieved in a large range of meristem sizes. Moreover, if *n ≈* 2*N*_b_ or larger, then the optimal probability distribution of apical stem cell divisions is close to uniform for the first *≈* 2*N*_b_ levels (and zero afterwards). This is very easy to implement by living systems, because a uniform pattern of cell divisions requires the least control (as the cells need not be distinguished in terms of their divisional rates).

The fact that the theoretical minimum of the average branching cost increases (although only logarithmically) with the average number of buds per branching (*N*_b_), explains why plants concentrate their branching capability to discrete locations (buds), and do not distribute it homogeneously.

Our results are in line with experimental studies conducted by Burian et al. [11], which found that the cells from which the axillary meristems originate, arise from the boundary of the apical meristems zone. Their results imply that mutational load in plants is primarily determined by branching order, and not by the length of branches [11, 13], which is also in agreement with the mechanism described above. Because the branching cost increases very slowly (only logarithmically) with increasing *N*_b_, the rate of molecular evolution is expected to be lower in plants with longer generation times, as plant longevity is primarily proportional to the branching order. This is consistent with the observation that rates of molecular evolution in plants are inversely proportional to generation times [14].

In conclusion, we describe an effective stochastic mechanism by which stem cells of plant meristems are able to maintain their genetic fidelity for long times, while producing a large number of branches and an even larger number of buds, each with the potential of initiating a new branch. In general, our results suggest that any branching system (not only plants), the growth of which relies on the duplication of its building blocks (cells), and in which a branching potential needs to be maintained all along its entire structure, should employ the mechanism described above to minimize the accumulation of copy errors.

## METHODS

### Numerical method

For any given values of *n* and *p_i_* Eq. (3) was solved numerically, and the numerical results for *D_n_*(*t*) were plugged into Eq. (5). Because Eq. (5) contains an infinite sum, it has been truncated at a large enough time value, and the rest of the infinite some approximated as follows.

After the transient time

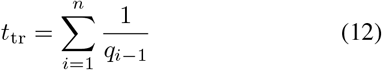

all the *D_i_*(*t*) values converge to a linear growth with the rate of *p*_0_, because all the cells of the meristem become the descendants of the cell at level 0. Thus, for *t ≫ t*_tr_ and for *t*^I^ *>* 0

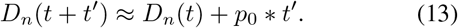

Using this relationship, for a large but finite truncation time *T ≫ t*_tr_ Eq. (5) can be substituted by the approximative formula

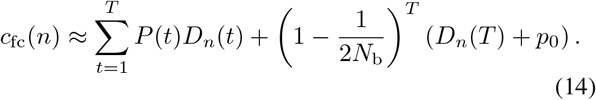

### Numerical optimization

For given values of *N*_b_ and *n* we initially set a uniform probability distribution *p_i_* = 1*/n* and determined *c*_fc_(*n*) using the above approximative formula. Then, after each step of the optimization we relocated a small random fraction of a randomly selected *p_i_* to one of its neighbors and determined the new value of *c*_fc_(*n*). We accepted the move if the new value was smaller than the value before the move. We stopped the optimization when the frequency of the accepted moves dropped below a small predefined threshold.

## References

[1] S. Smith, F1000 - Post-publication peer review of the biomedical literature (2017), 10.3410/f.726838551.793530056.

[2] R. Lanner, Ageing Research Reviews 1, 653671 (2002).

[3] “Rocky mountain tree-ring research,”.

[4] S. C. Sillett and R. V. Pelt, Ecological Monographs 77, 335359 (2007).

[5] W. D. Flint, To find the biggest tree (Sequoia Natural History Association, 2002).

[6] R. Lanfear, PLOS Biology 16(2018), 10.1371/journal.pbio.2005439.

[7] L. Wang, Y. Ji, Y. Hu, H. Hu, X. Jia, M. Jiang, X. Zhang, L. Zhao, Y. Zhang, Y. Jia, and et al., PLOS Biology 17(2019), 10.1371/journal.pbio.3000191.

[8] C. Plomion, J.-M. Aury, J. Amselem, T. Leroy, F. Murat, S. Duplessis, S. Faye, N. Francillonne, K. Labadie, G. L. Provost, and et al., Nature Plants 4, 440452 (2018).

[9] D. Francis, Encyclopedia of Life Sciences (2008), 10.1002/9780470015902.a0002050.pub2.

[10] E. Schmid-Siegert, N. Sarkar, C. Iseli, S. Calderon, C. Gouhier-Darimont, J. Chrast, P. Cattaneo, F. Schütz, L. Farinelli, M. Pagni, et al., Nature Plants 3, 926 (2017).

[11] A. Burian, P. Barbier De Reuille, and C. Kuhlemeier, Current Biology 26, 13851394 (2016).

[12] I. Derényi and G. J. Szöllősi, Nature Communications 8, 14545 (2017).

[13] E. P. Groot and T. Laux, Current Biology 26(2016), 10.1016/j.cub.2016.05.049.

[14] A. R. D. L. Torre, Z. Li, Y. V. D. Peer, and P. K. Ingvarsson, Molecular Biology and Evolution 34, 13631377 (2017).

